# A simple growth model for tumor spheroids

**DOI:** 10.1101/2025.10.02.680182

**Authors:** Uwe Schneider, Ning Liang

## Abstract

Mathematical models of tumor spheroid growth provide essential insights into avascular tumor dynamics, hypoxia development, and treatment response. Here, we present a simple biophysically motivated growth model that integrates energy conservation, oxygen diffusion, and necrosis formation into a unified framework. The model captures the key features of spheroid development, including initial exponential growth, deceleration due to diffusion limitation, and growth inhibition mediated by necrotic tissue. Using published growth data from human colon carcinoma (HCC1), mouse colon carcinoma (MCC26), V79 fibroblasts, and EMT6-Ro spheroids, we demonstrate that the model accurately reproduces observed growth kinetics across multiple phases. Importantly, the model predicts viable rim thickness and oxygen distribution over time, in agreement with theoretical expectations. Fitted parameters further enable estimation of single-cell mass, oxygen consumption rate, and the energetic cost of cell production. By linking geometry, metabolism, and growth regulation, this minimal model offers a transparent yet quantitative description of tumor spheroid behavior, with potential applications in radiobiology and therapy modeling.

## Introduction

Multicellular tumor spheroids are widely used as an *in vitro* model of avascular tumor growth. They replicate key features of early tumor development, including three-dimensional architecture, gradients of oxygen and nutrients, and the emergence of central necrosis. Because of their reproducibility and experimental accessibility, spheroids have become a standard model system in radiobiology, pharmacology, and tumor physiology research.

Several mathematical models have been proposed to describe spheroid growth, ranging from purely empirical fits to detailed biophysical simulations. Classical Gompertzian or logistic models reproduce the sigmoidal growth curve but do not explicitly account for oxygen limitation or necrosis formation. More mechanistic approaches incorporate diffusion and consumption of oxygen and nutrients, predicting the formation of a necrotic core once the spheroid exceeds a critical size. Despite these advances, many existing models are either too abstract to provide mechanistic insight or too complex for routine application. There remains a need for simple yet biophysically grounded models that can link fundamental energy constraints to measurable growth dynamics. In this work, we develop and test such a model. Building on conservation of energy and oxygen diffusion, we derive a three-phase description of spheroid growth: (i) initial exponential expansion, (ii) growth slowdown due to diffusion limitation and necrosis onset, and (iii) further inhibition mediated by necrotic factors.

We validate the model using published data on human colon carcinoma (HCC1), mouse colon carcinoma (MCC26), V79 fibroblast, and EMT6-Ro spheroids. The model reproduces growth curves, viable rim thickness, and scaling relations across a wide range of spheroid sizes. In addition, parameter estimation yields biologically meaningful quantities such as average cell mass, oxygen consumption rate, and the energetic cost of cell production. Our approach balances simplicity and interpretability, providing a transparent framework for analyzing spheroid growth and its implications for tumor biology and therapy.

## Materials and Methods

### Tumor spheroid growth equation

The growth of tumor spheroids is governed by the balance between nutrient and oxygen supply and cellular demand. Initially, spheroids develop from single cells and rely solely on the diffusion of oxygen and nutrients across their outer surface to sustain proliferation. As the spheroid enlarges, diffusion becomes insufficient to support cells in the inner regions, leading to the formation of a necrotic core. Continued growth is then influenced by both structural changes and biochemical feedback mechanisms, including the presence of necrotic tissue. Previous studies (e.g., Freyer [4]) suggest that tumor necrosis factor or other inhibitory substances derived from necrotic or dying cells may suppress the proliferation of viable cells. While the exact mechanism remains unclear, it is hypothesized that either quiescent cells transitioning to necrosis or the necrotic cells themselves may release diffusible factors that inhibit further growth. This regulatory feedback plays a critical role in limiting spheroid expansion and is accounted for in the modeling framework presented here. Conservation of energy requires that the total metabolic rate of a tumor spheroid, *B_T_*, be apportioned between the power required for maintenance and that for mitosis [3]:

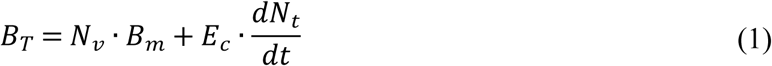

where *N_v_* is the number of viable cells at time *t* after growth begins, *N_t_* the total number of cells, *B_m_* the power each cell requires for maintenance, and *E_c_* the energy to create a cell. We follow here the argumentation of Herman et al. [3], who assume that the rate of increase of the number of viable cells is the difference between the total rate of mitosis (*dN_t_/dt*) and the rate of cell death due to apoptosis and necrosis (*dN_n_/dt*). Moreover, independent of the specific type of cell death, an average life time (1/Γ) of a viable cell can be defined. Thus Eq.1 yields:

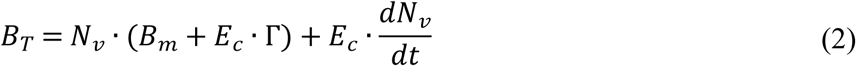

If we now define the average cellular metabolic power required for maintenance and replacement *B_c_ = B_m_ + E_c_·Γ* and *m_c_* as the average mass of a cell, Eq.2 transforms into a differential equation for the viable mass *m_v_*:

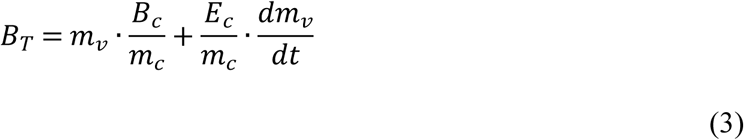

Up to this point, the derivation of this differential equation describing the conservation of energy does not yet differ from Herman et al [3] for describing tumors. However, solving this equation requires knowledge of how tumor metabolic rate, *B_T_*, depends on its viable mass, *m_v_*, and this will need another approach for tumor spheroids than for tumors. But let’s turn first to the oxygen distribution in spheroids.

### The oxygen distribution in tumor spheroids

Understanding oxygen distribution in tumor spheroids is essential for predicting regions of hypoxia, cellular viability, and treatment response. Due to the lack of vasculature in avascular tumor spheroids, oxygen is supplied solely by diffusion from the surrounding medium and is consumed by cells throughout the spheroid. As a result, a gradient forms, often leading to the development of a anoxic core at sufficient spheroid sizes.

In this work, we adopt the framework developed by Grimes et al. [5], which provides a method to estimate the oxygen partial pressure in multicellular tumor spheroids based on the steady-state diffusion equation. This approach is grounded in the solution of Fick’s second law under spherical symmetry and considers uniform oxygen consumption within the viable rim of the spheroid. By assuming steady-state diffusion and neglecting convection, the oxygen partial pressure profile *p(r)* as a function of radius *r* is:

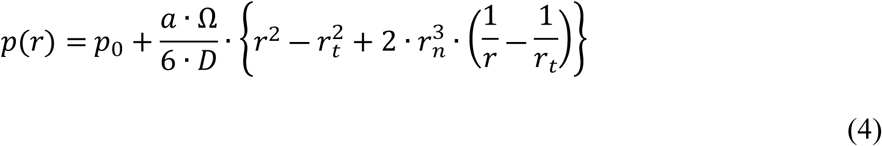

where *p_0_* is the oxygen partial pressure at the surface of the tumor spheroid, *a* is the oxygen consumption rate, and *D* is the diffusion coefficient of oxygen in the tissue. For a spheroid of radius *r_t_*, the solution to this equation allows for the determination of the oxygen distribution throughout the spheroid, including the critical radius *r_n_* at which the oxygen level drops to zero at the center, indicating the onset of central necrosis. This is illustrated in figure 1 with the thickness of the viable rim being *r_v_* = *r_t_ - r_n_*.

**Figure 1.**
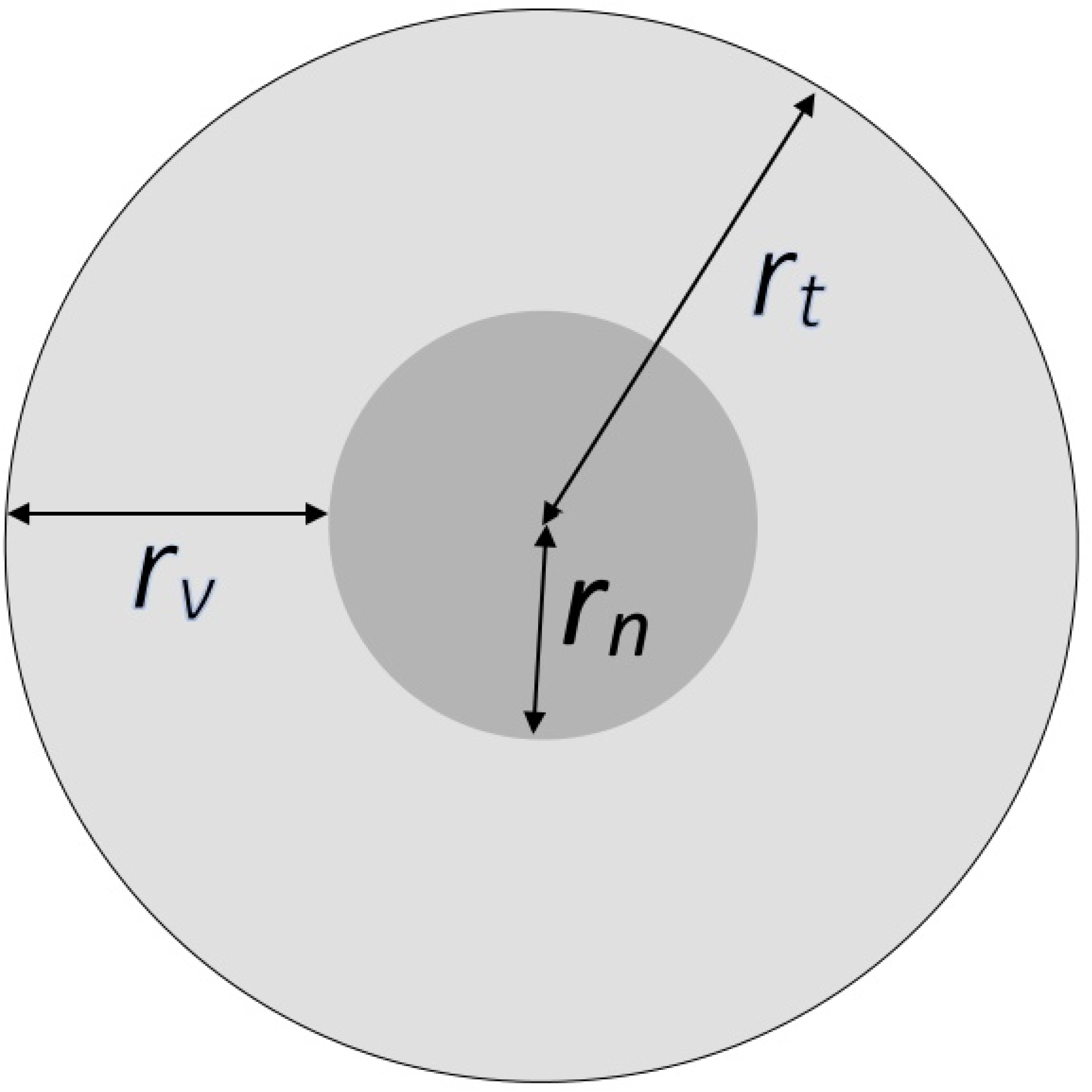
Schematic cross-section of a tumor spheroid of radius *rt*. The viable region, of thickness *rv*, surrounds the necrotic core (*r_n_*) where oxygen is absent. The oxygen distribution determines the viable rim and onset of central necrosis.

One of the key insights from Grimes et al. [5] is the use of experimentally measurable parameters such as spheroid radius and external oxygen concentration *p_0_* - to estimate *a* indirectly. Their method relies on detecting the threshold at which necrosis initiates, corresponding to the radius *r_l_* where oxygen reaches zero at the core:

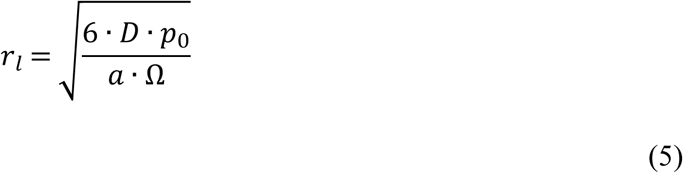

where the oxygen diffusion constant *D* is close to that of water (*D* = 2·10^-9^ m^2^ s^-1^), Ω is a constant (Ω = 3.0318·10^7^ Torr kg^-1^m^-3^) and *a* is the oxygen consumption rate in m^3^ kg^-1^ s^-1^. The conversion of *a* into cellular metabolic power of a cell is given in Appendix A. Grimes et al. [5] provide also a general solution for the necrotic core radius:

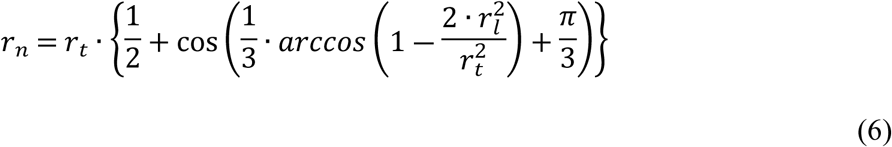

And if we assume that the density of tumor spheroids is *ρ* = 1 g/cm^3^ and always perfectly spherical in geometry, masses can be directly calculated from the radii: *m* = 4/3ˑπˑr^3^ with the mass given in g and the radius in cm. The viable mass of the tumor spheroid is then simply:

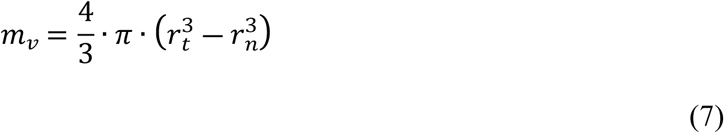

And this yields to the very important result of a constant rim thickness for increasing spheroid size:

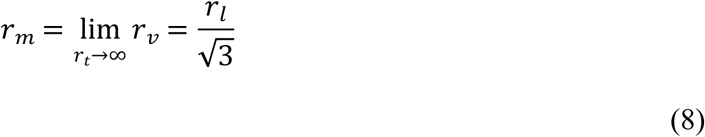

Incorporating this model into the broader tumor growth framework enables time-resolved simulation of oxygen profiles during spheroid expansion. As the spheroid grows and oxygen consumption rate remains relatively constant, the oxygen penetration depth decreases, eventually resulting in a hypoxic core. This distribution can be used to infer biologically important metrics such as the hypoxic fraction, viable rim thickness, and spatially resolved radiosensitivity—all of which play a crucial role in radiotherapy treatment planning and therapeutic resistance.

### The metabolic rate and mass scaling of tumor spheroids

#### Growth phase 0

In tumor spheroids, the cellular oxygen consumption rate, *a*, can be treated as approximately constant [5]. Consequently, the metabolic rate per tumor cell is given by

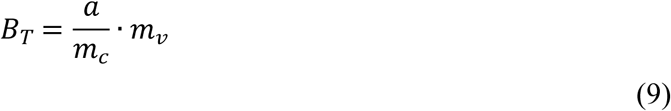

In the initial stage of tumor development, oxygen diffuses uniformly throughout the spheroid. Thus, the total tumor mass and the viable tumor mass are effectively identical, and equation (3) becomes:

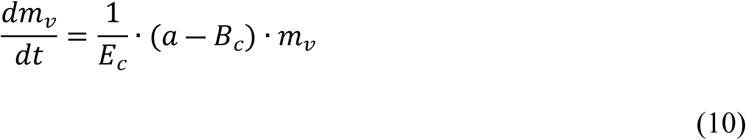

Equation (10) leads to exponential growth for both the total and viable tumor masses:

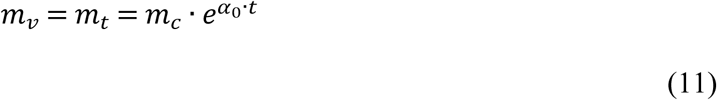

with

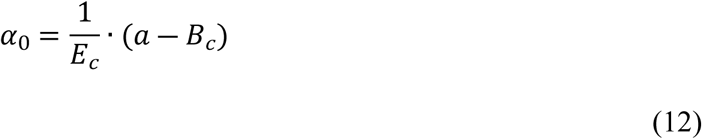

The qualitative behavior of the growth curve of tumor spheroids is sketched in Figure 2.

**Figure 2.**
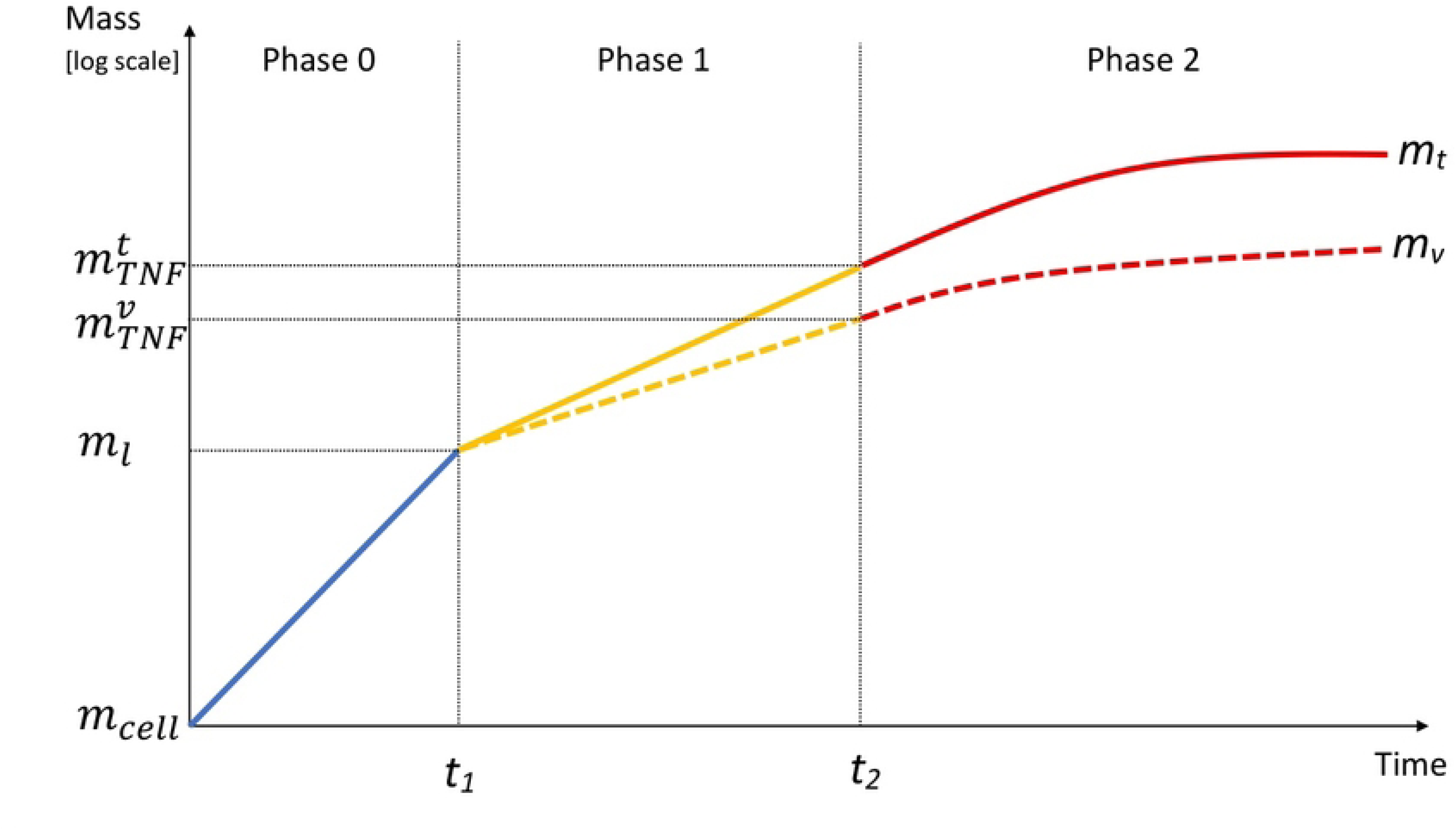
Qualitative growth curves for total and viable tumor spheroid mass during the three modeled growth phases. The onset of necrosis at *t_1_* and the transition to growth inhibition at *t_2_* are indicated.

#### Growth phase 1

In the early regime in which a necrotic core occurs (*t>t_1_* with *m_t_(t_1_) = m_l_*), our model still predicts a metabolic rate according to equation (9) and exponential growth continues for the viable mass. However, in this regime, viable and total tumor mass are now non-linearly related. This is because the viable mass forms a spherical shell around the necrotic core. Therefore, the viable mass is approximately proportional to the square of the total spheroid radius and it is:

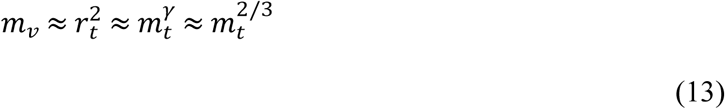

The exponent *γ* = 2/3 can also be derived using the Grimes formalism, applying equation (7) for *m_v_* in the limit of large *m_t_*. However, equation (7) shows that the exact value 2/3 is only attained for extremely large tumor spheroid masses. Experimental evidence [4] indicates that the ratio *m_t_/m_l_*, where *m_l_* is the mass at which the central oxygen pressure drops to zero, is at most 100. For this biologically relevant range, equation (7) is fitted in Appendix C. For phase 1 this yields:

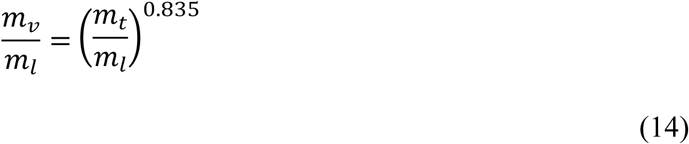

In this regime, the total tumor mass grows exponentially at a faster rate than the viable mass, leading to a steadily increasing fraction of necrotic tissue. This can be expressed as

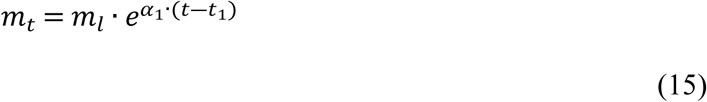

Thus, fitting equation (15) to experimental spheroid growth data automatically yields the viable mass through equation (14).

#### Growth phase 2

As the necrotic core expands, extracts from necrotic tumors have been shown to inhibit the growth of tumor spheroids [1, 4], although the underlying mechanism remains unclear. One possibility is that necrotic cells release a factor that suppresses the proliferation of viable cells [1]. Consequently, the release of this tumor necrosis factor (TNF) is expected to scale with the necrotic mass. From equation (8), we know that for sufficiently large tumor spheroids the thickness of the viable rim remains constant. Thus, when *r_n_ ≈ r_t_*, the necrotic mass becomes directly proportional to the total spheroid mass. Moreover, since at larger total masses we have 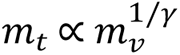 (with *γ* = 0.713 from Appendix C), an additional loss term representing the effect of TNF on viable mass can be incorporated into equation (9) to describe phase 2 growth dynamics:

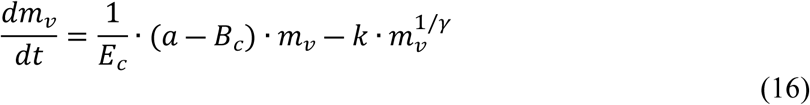

- If we define 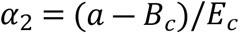 and write 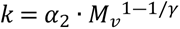, where *M_v_* is the asymptotic viable mass of the tumor, the general solution of equation (17) is

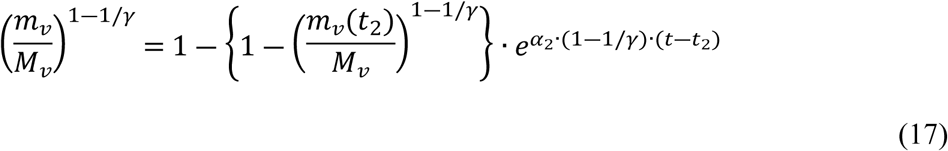

where *m_v_(t_2_)* is the viable malignant mass at the transition time *t_2_* which can be calculated from equations (14) and (15). Accordingly, we can calculate the total mass in growth phase 2, where *M_t_* is the asymptotic total mass of the tumor:

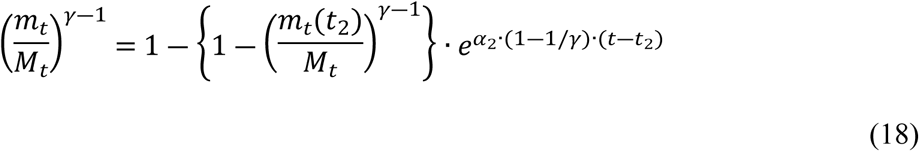

Since the cellular oxygen consumption rate *a* remains approximately constant during tumor spheroid growth [5], it can be determined from the limiting radius *r_l_* and the single-cell mass *m_c_*, both obtained from the data fit. From equation (12), it follows that the energy required to generate a new cell satisfies *E_c_*> *a/α*. This allows to estimate an upper limit for the energy to create a cell.

### Experimental tumor spheroid growth data and model fit

Model verification was performed using experimental tumor spheroid growth data spanning a wide range of spheroid volumes. Specifically, we used human colon carcinoma (HCC1) and mouse colon carcinoma (MCC26) data reported by Freyer [4], together with V79 fibroblast spheroid data from Marušić et al. [4, 8]. Both studies also provide measurements of viable rim radii, which, in conjunction with equation (8), allow determination of the limiting radius *r_l_*, the corresponding spheroid mass *m_l_*, and the associated time *t_1_* at which diffusion limitation occurs.

Equation (11) was then fitted to the data by least-squares minimization, yielding estimates of the single-cell mass *m_cell_* and the parameter *α_0_*. The transition time *t_2_*, marking the onset of the production of tumor necrosis factor, is not experimentally available. Therefore, *t_2_*was fitted simultaneously with *α_1_* and *α_2_* under the constraint that the slope of the model in growth phase 1 (equation (15)) coincides with the slope of the model in growth phase 2 (equation (18)) at *t_2_*.

## Results

### Model fitting to experimental spheroid growth

The model was validated using experimental growth data from human colon carcinoma (HCC1), mouse colon carcinoma (MCC26) [4], V79 fibroblasts [8], and EMT6-Ro cells [9]. These datasets span a wide range of spheroid sizes and growth kinetics, thereby providing a robust test of model performance. For each dataset, the total spheroid mass *m_t_* was fitted and the viable mass *m_v_* computed from the model. Transition times *t_1_* (onset of necrosis) and *t_2_* (onset of necrotic inhibition) were estimated by least-squares minimization, with the constraint that growth phase 1 and phase 2 slopes coincide at *t_2_*.

The model accurately reproduced the experimental growth curves across all four cell lines (Figure 3). The three distinct growth phases were clearly resolved: (i) initial exponential increase, (ii) reduced growth following oxygen limitation and necrosis onset, and (iii) further slowdown due to inhibitory effects of necrotic tissue. The agreement between predicted and measured data supports the validity of the framework across different tumor types.

**Figure 3.**
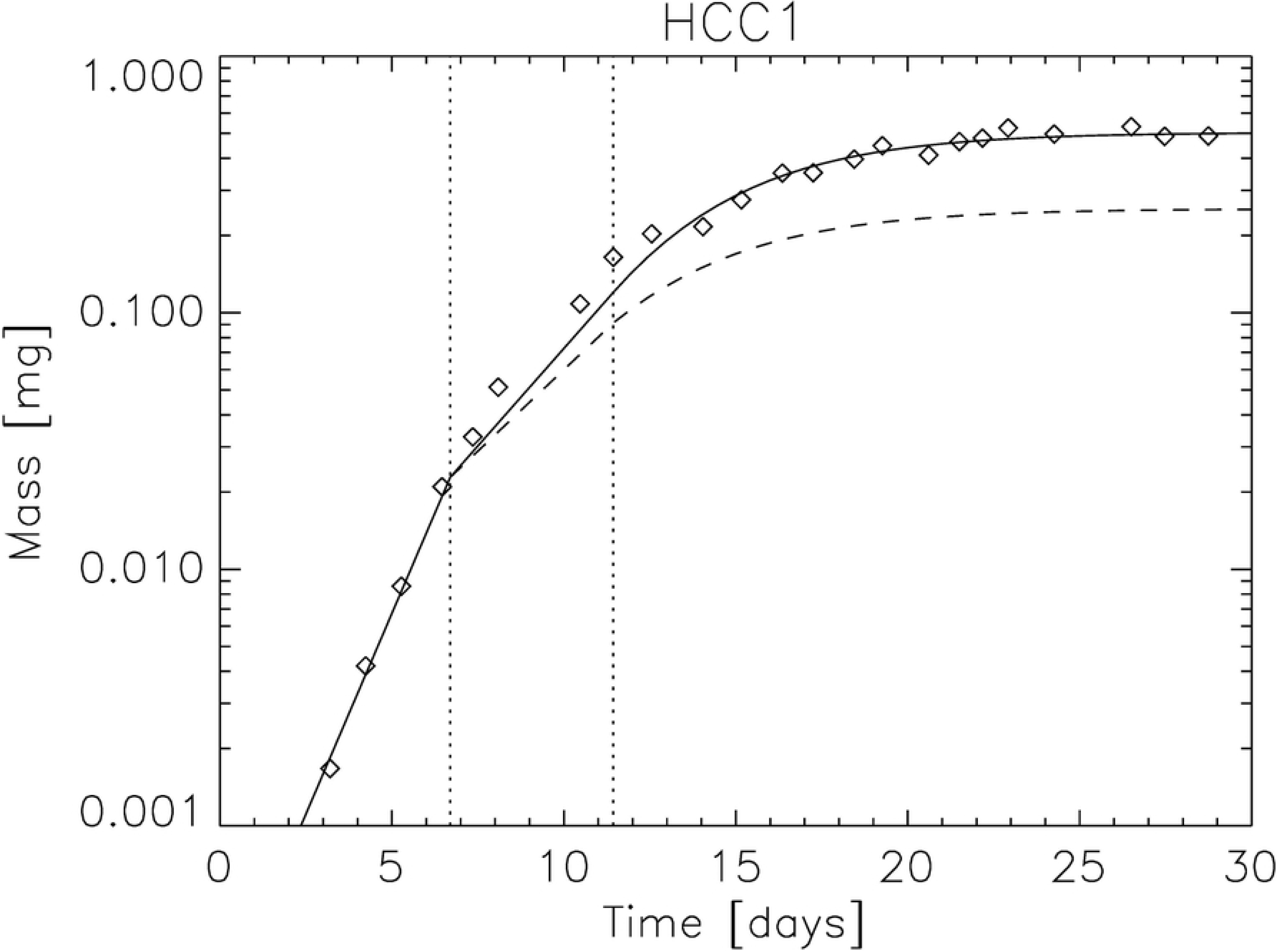
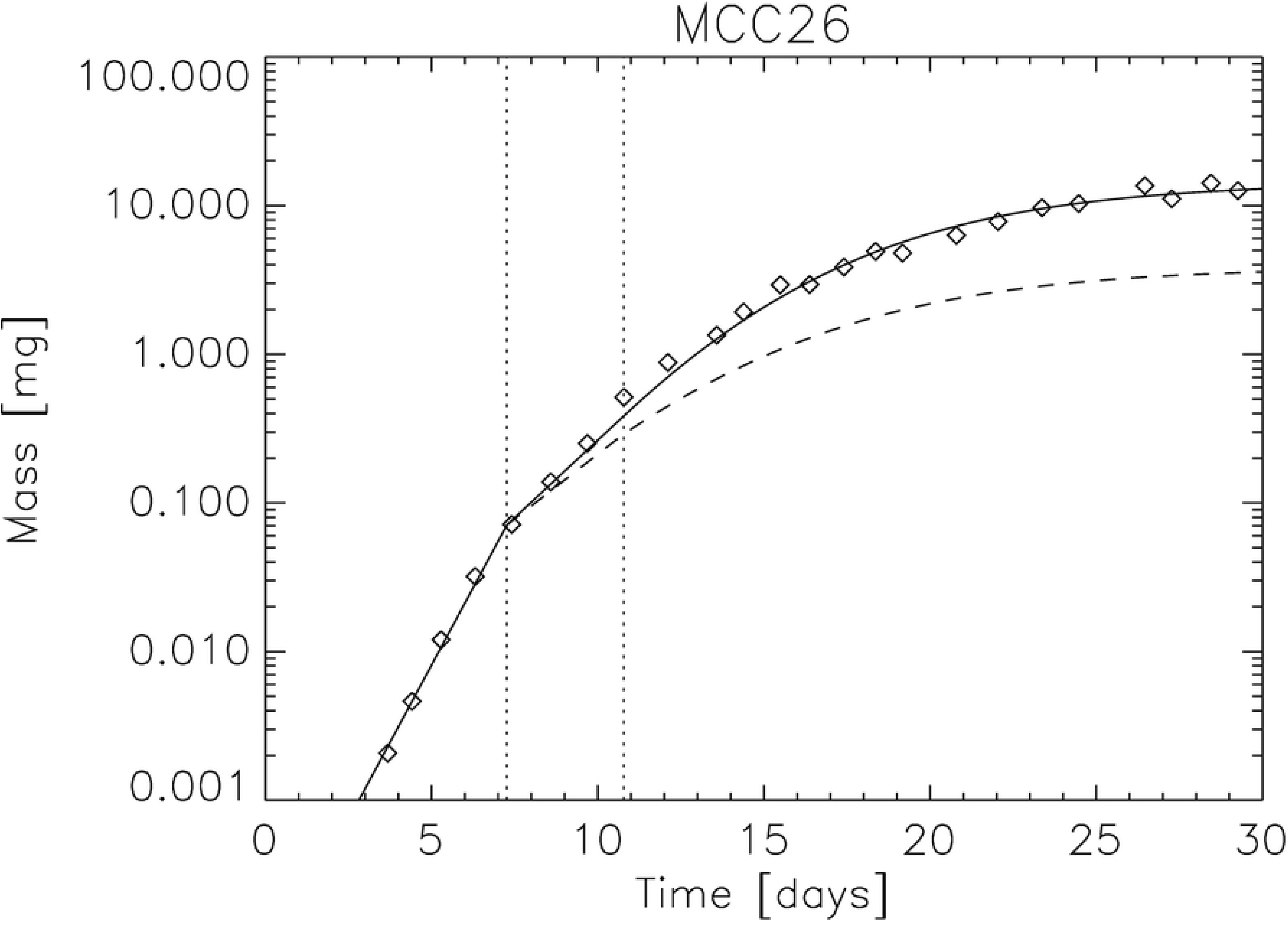
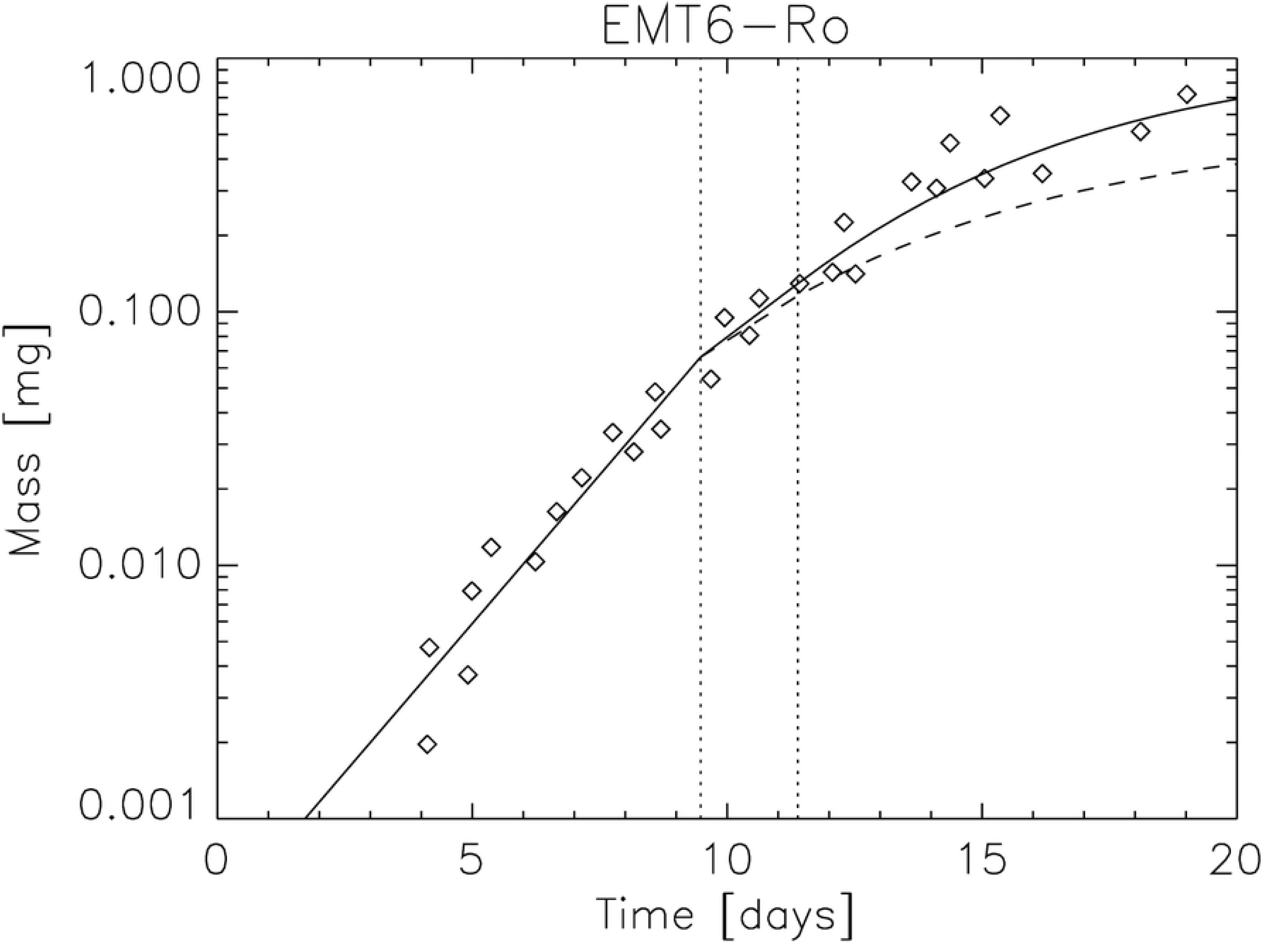
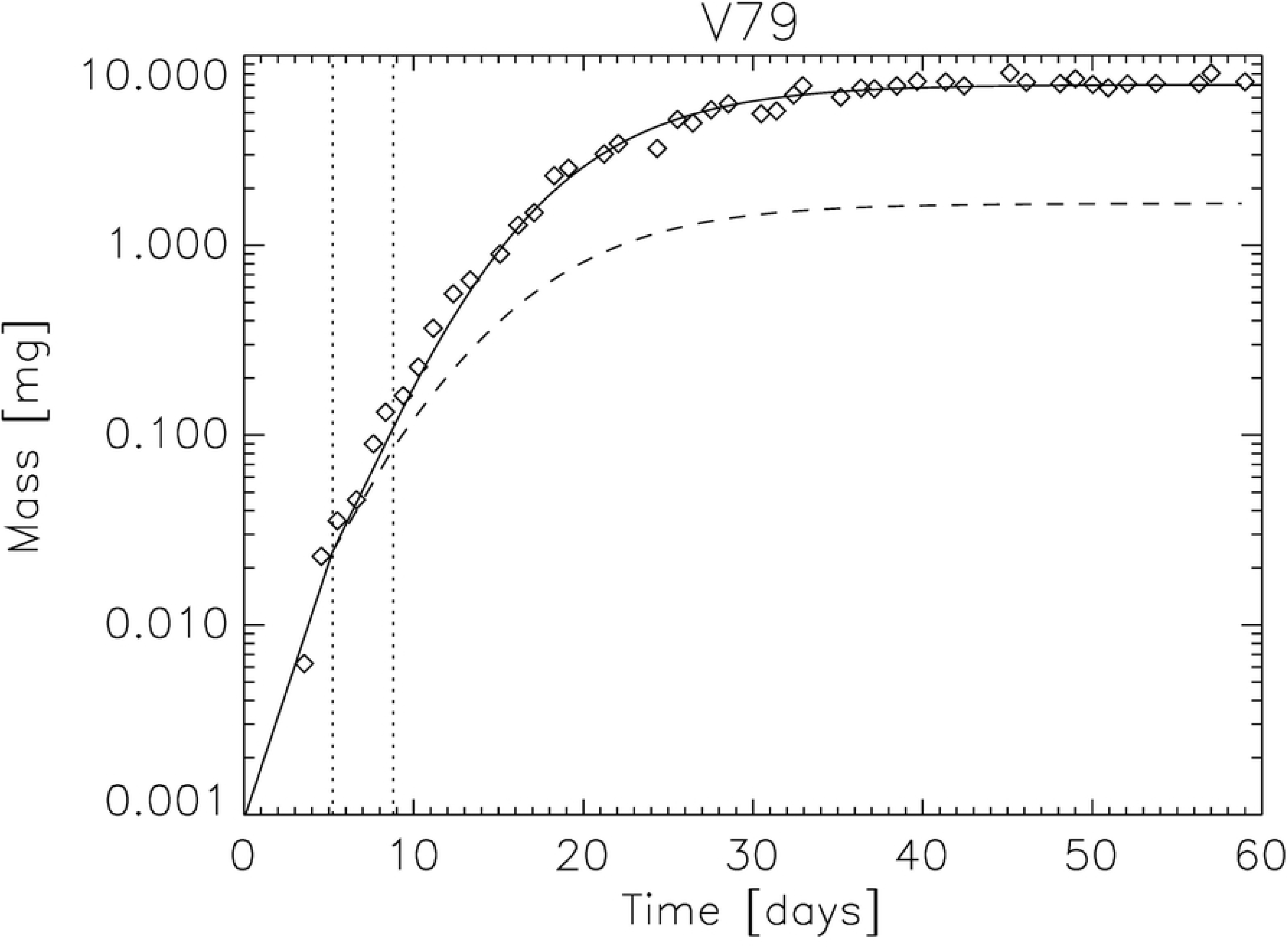
Fitted growth curves of total tumor spheroid mass (solid lines) for four cell lines: HCC1, MCC26, V79, and EMT6-Ro. Dashed lines indicate corresponding viable mass. Vertical dotted lines mark transition times *t_1_* and *t_2_*.

### Estimation of cell mass and oxygen consumption

From the fitted parameters, the average single-cell mass *m_c_* and oxygen consumption rate *a* were estimated (Table 1). For all four datasets, values of *m_c_* were consistent with previously reported ranges for mammalian cells, typically on the order of 10^-14^ - 10^-15^ kg. The corresponding oxygen consumption rates per cell were also within the expected physiological range and similar to values derived independently from spheroid oxygen diffusion studies [5].

**Table 1.**

| Parameter |  | Cell type |  |  |  |
| --- | --- | --- | --- | --- | --- |
|  |  | HCC1 | MCC26 | V79 | EMT6-Ro |
| $m_{\text{cell}}$ | [kg] | 1.84E-13 | 6.76E-14 | 9.47E-13 | 3.95E-13 |
| $\alpha_0$ | [s<sup>-1</sup>] | 0.72 | 0.96 | 0.62 | 0.54 |
| $T_1$ | [s] | 6.7 | 7.3 | 5.2 | 9.5 |
| $\alpha_1$ | [s<sup>-1</sup>] | 0.35 | 0.48 | 0.42 | 0.35 |
| $T_2$ | [s] | 11.4 | 10.8 | 8.8 | 11.4 |
| $\alpha_2$ | [s<sup>-1</sup>] | 0.74 | 0.53 | 0.43 | 0.56 |
| $M_t$ | [kg] | 5.04E-10 | 1.44E-8 | 7.01E-9 | 1.01E-9 |
| $P_0$ | [Torr] | 150 | 150 | 150 | 150 |
| $r_m$ | [m] | 1.76E-4 | 2.57E-4 | 1.80E-4 | 2.51E-4 |
| $r_l$ | [m] | 3.05E-4 | 4.45E-4 | 3.12E-4 | 4.35E-4 |
| $a$ | [m<sup>3</sup>·kg<sup>-1</sup>·s<sup>-1</sup>] | 1.92E-6 | 8.99E-7 | 1.83E-6 | 9.42E-7 |
| $a$ | [J·s<sup>-1</sup>] | 2.12E-12 | 3.65E-13 | 1.04E-11 | 2.23E-12 |
| $E_c(\text{max})$ | [J] | 6.05E-12 | 7.60E-13 | 2.48E-11 | 6.38E-12 |

### Scaling of viable mass with total mass

A key prediction of the model is the scaling relation between viable and total mass. For large spheroids, purely geometric considerations suggest 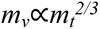. To test this, experimental data were compared with both the Grimes model [5] and a two-phase power-law fit of the form ln(*m_v_*/*m_l_) = γ·*ln(*m_t_*/*m_l_) +* ln(*m_v_*(0)/*m_l_).* The fit yielded exponents *γ* = 0.835 in the early diffusion-limited regime (Phase 1) and *γ* = 0.713 at larger sizes (Phase 2), in excellent agreement with model predictions (Figure C1). The maximum deviation between the fitted curve and Grimes’ theoretical data was less than 0.2%, confirming the robustness of the approximation.

### Prediction of viable rim thickness

The model also predicts the viable rim radius *r_v_* as a function of total spheroid radius *r_t_*. As expected, the viable rim approached a constant thickness at large *r_t_*, consistent with oxygen diffusion theory. Predicted rim sizes were in close agreement with experimental measurements and converged towards the theoretical limit reported by Grimes et al. [5] (Figure 4). Minor deviations at intermediate sizes reflect cell line–specific differences in necrosis onset and growth inhibition.

**Figure 4.**
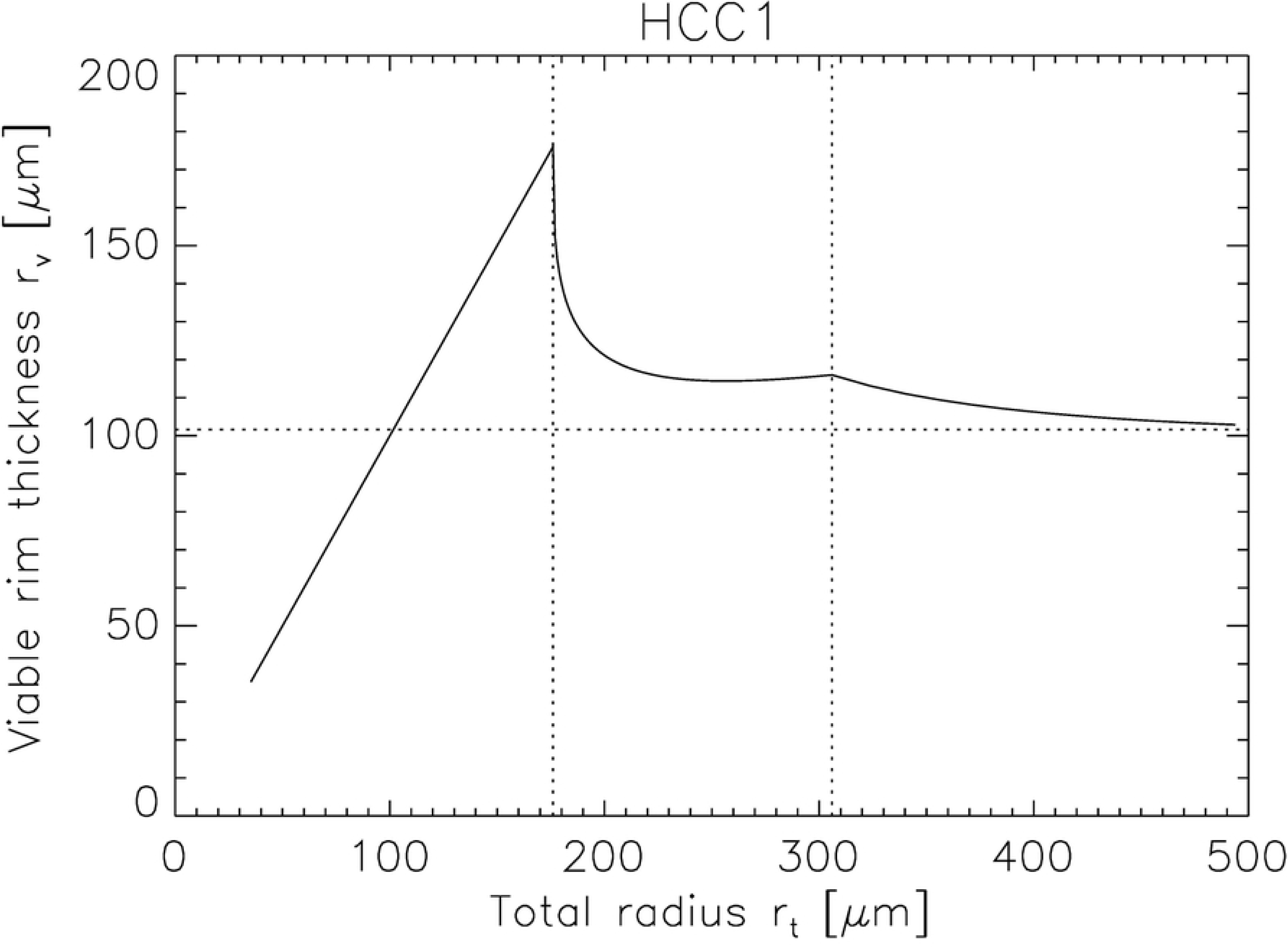
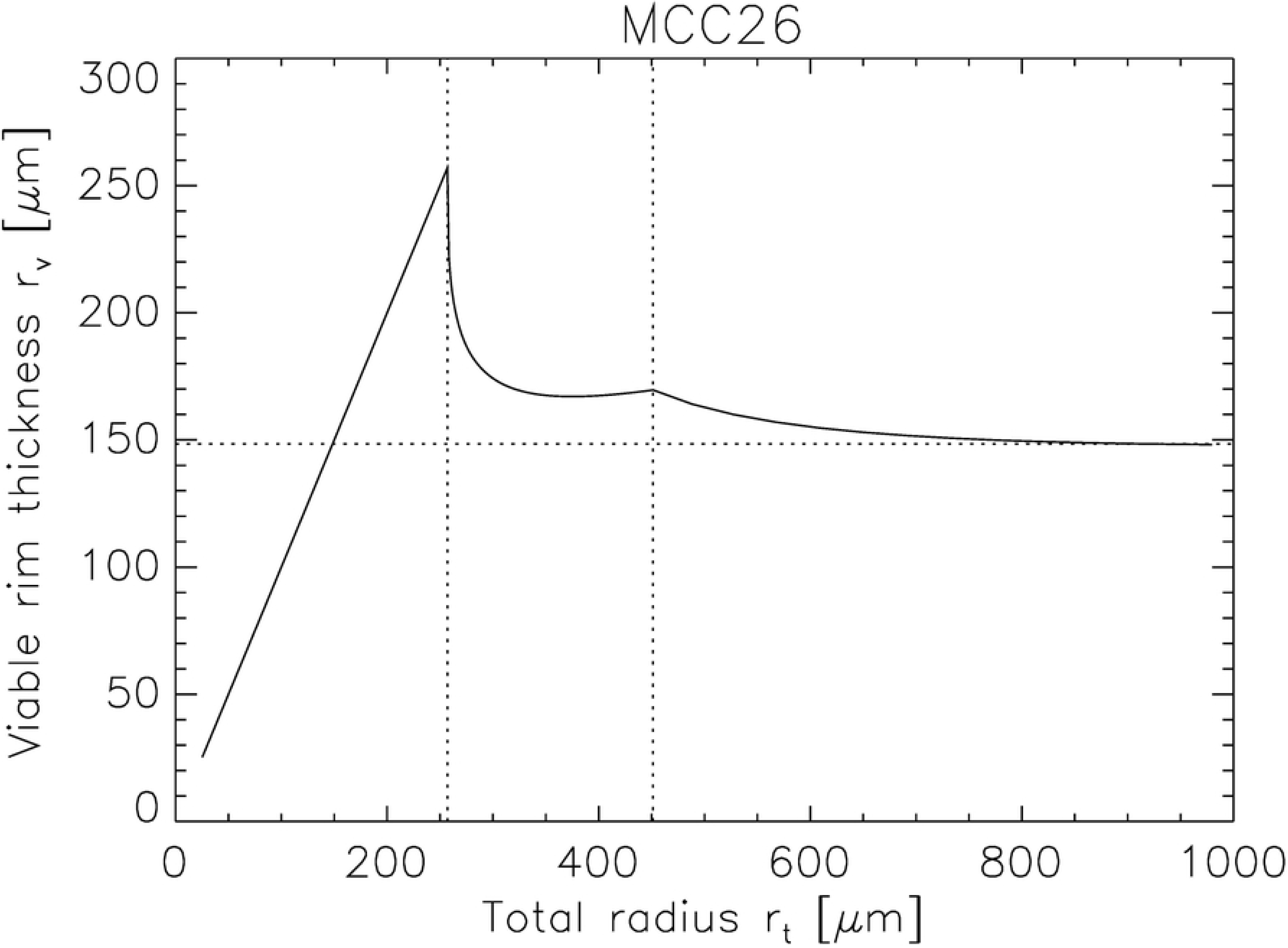
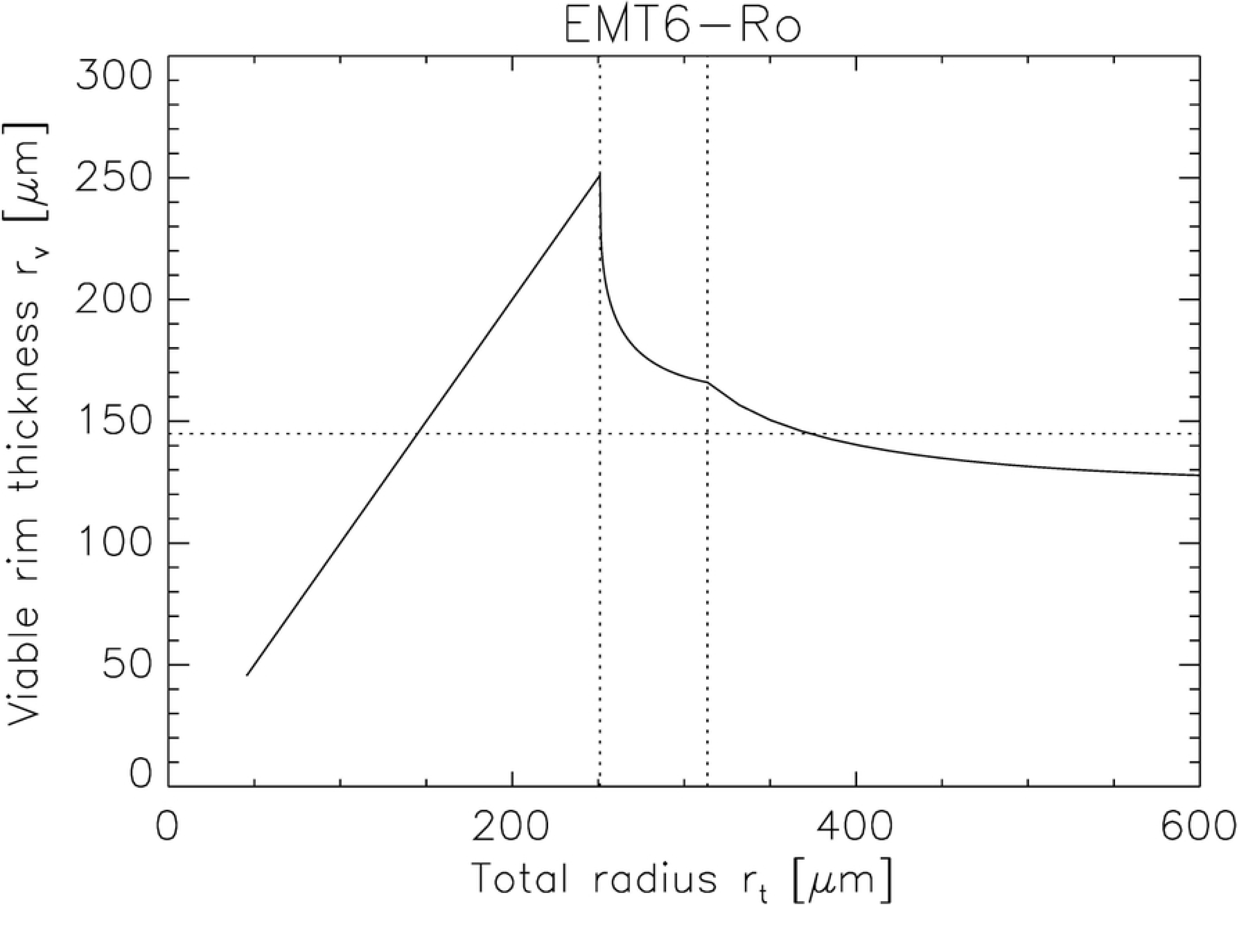
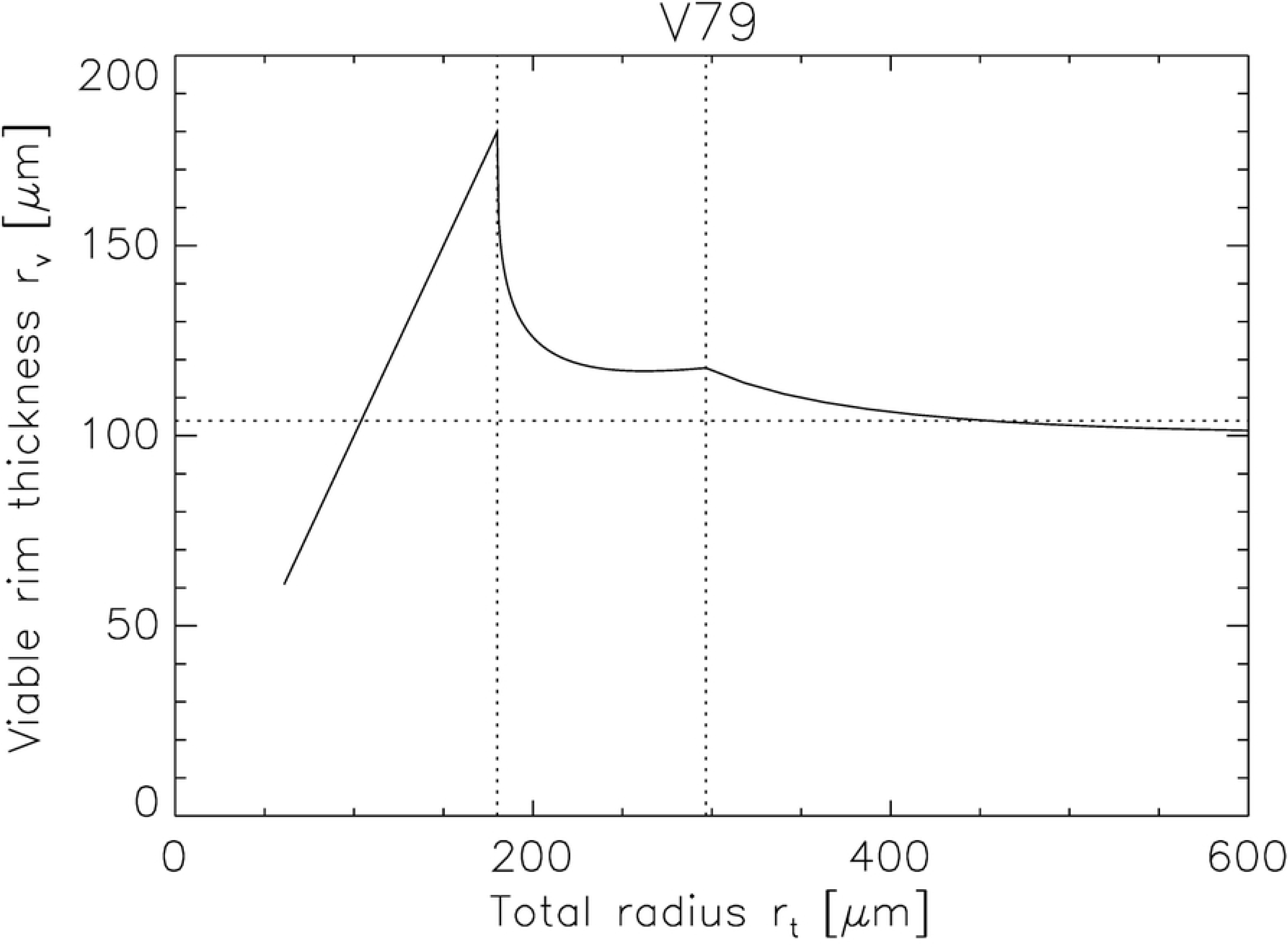
Predicted viable rim thickness as a function of total spheroid radius for four cell lines. Vertical dotted lines indicate transition times *t_1_* and *t_2_*. The horizontal dotted line shows the theoretical limit of viable rim size predicted by Grimes et al. [5].

### Energetic cost of cell production

Finally, the model provides an estimate of the energy required to generate a new cell. Based on equation (12), the energy per cell is bounded by *E_c_* > a/α. For the datasets analyzed here, fitted parameters indicate that the energetic cost of cell production is on the order of a few picomoles of ATP per cell, in line with independent biochemical estimates. Importantly, this partitioning of total energy consumption into maintenance and growth terms provides a quantitative framework for evaluating how metabolic constraints shape tumor expansion.

## Discussion

Our study introduces a mechanistically grounded but computationally simple model that successfully captures the main features of tumor spheroid growth across multiple cell types. The model builds upon energy conservation principles and explicitly accounts for oxygen diffusion, necrosis formation, and growth inhibition by necrotic tissue. By doing so, it provides a direct connection between biophysical constraints and macroscopic growth behavior.

A key result is the model’s ability to predict the scaling of viable mass with total spheroid mass. While geometric considerations suggest an asymptotic exponent of 2/3, fits to biologically relevant data yield higher effective exponents for small spheroids (γ ≈ 0.83) and somewhat lower values at larger sizes (γ ≈ 0.71). These findings are consistent with experimental reports that oxygen limitation alone cannot fully explain growth saturation, and that necrotic cell–derived inhibitory factors contribute substantially in later phases.

The fitted parameters provide biologically meaningful quantities, including estimates of average cell mass, oxygen consumption rate, and the maximum energy cost of cell production. These values are in line with previously reported ranges for mammalian cells, supporting the plausibility of the framework. Importantly, the model also provides a quantitative description of viable rim thickness, which converges towards theoretical predictions by Grimes et al. [5]. Deviations at intermediate sizes highlight the biological complexity of necrosis induction and growth inhibition, which may differ between cell lines.

Beyond growth kinetics, the explicit coupling between oxygen distribution and viable cell mass has important implications for radiobiology. Hypoxia is a key determinant of radiosensitivity, and the ability to predict its spatial and temporal dynamics from simple geometric and metabolic parameters opens opportunities for integrating growth models into radiotherapy planning. Although demonstrated here for a limited number of cell lines, the generality of the model suggests broad applicability to spheroids and early avascular tumors.

Limitations of this work include the assumption of constant oxygen consumption per cell, uniform cell density, and simplified treatment of necrotic inhibition. Incorporating variable metabolic states, cell cycle heterogeneity, and detailed biochemical signaling would refine predictions at the cost of simplicity. Nevertheless, our approach balances parsimony and interpretability, offering a transparent framework that can be readily applied and extended.

## Conclusion

A central advantage of this approach lies in its simplicity. Whereas many existing models rely on systems of coupled partial differential equations or complex parameterizations of nutrient transport and cellular metabolism, our framework reduces the problem to a small set of interpretable equations. The resulting parameters—cell mass, oxygen consumption rate, and energetic cost of cell production—are directly linked to measurable biological quantities. This parsimony allows the model to be applied and fitted to experimental data with minimal computational overhead, while retaining a clear mechanistic foundation.

The model is therefore substantially simpler than most detailed continuum or agent-based approaches, yet it retains predictive power at the level of viable mass scaling and necrotic core dynamics. Its transparency makes it well suited for exploratory studies, integration into larger treatment models, and use as a baseline for more elaborate extensions that include variable metabolism, cell cycle effects, or microenvironmental interactions.

In summary, our model provides a tractable and generalizable framework for analyzing tumor spheroid growth. By striking a balance between simplicity and mechanistic insight, it offers a valuable tool for both experimental interpretation and theoretical investigation of avascular tumor dynamics.

## Competing Interests Statement

The authors state that there is no conflict of interest for the authors or the author’s institution and that they have no financial or personal relationships that inappropriately influence their actions. They have no dual commitments, competing interests, competing loyalties, employment, consultancies, stock ownership, honoraria, or paid expert testimony.

## Authors’ contributions

US had the idea for the project and developed the mathematical framework. NL contributed with results of her Master thesis. All authors read and approved the final manuscript.

## Appendix A: Conversion of Volumetric Oxygen Consumption to Cellular Metabolic Power

Grimes et al. report the oxygen consumption rate of tumor spheroids a in volumetric units of [m^3^ O2 kg^−1^ s^−1^]. To relate this to the metabolic power of an individual cell, the following conversion steps are required.

*Conversion from volume of O₂ to moles of O₂*

At physiological temperature (37 °C, 1 atm), the molar gas volume is approximately *V_m_* = 2.54×10^−2^ m^3^ mol^−1^. Thus, the molar consumption rate per unit tissue mass is *a_m_* = *a/V_m_* in units of [mol O_2_ kg^−1^ s^−1^].

*Conversion from moles of O₂ to metabolic power*

The complete reduction of one mole of O₂ via oxidative phosphorylation yields approximately *E_O2_* ≈ 1.525×10^5^ J mol^−1^, corresponding to five ATP per O atom at ΔG ≈ 30.5 kJ/mol ATP. The metabolic power per unit tissue mass is therefore *a_kg_* = *a_m_⋅E_O2_* in units of [J s^−1^ kg^−1^].

*Per-cell metabolic power*

To convert further the metabolic power per kg into the metabolic power per cell *a_c_*, the mass of a viable cell *m_c_*must be known. The average cell mass *m_c_* can be obtained by fitting the tumor spheroid growth model in phase 0 described by equation (11) to experimental data and is listed in Table (XX). The metabolic rate per cell is obtained by normalizing by this factor:

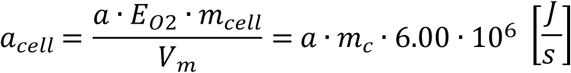

with *m_cell_* in [kg].

## Appendix B: Scaling of tumor viable mass as a function of the total tumor mass

The viable fraction of a tumor spheroid is localized in a spherical shell surrounding the necrotic core. Consequently, for large spheroids the viable mass scales approximately with the surface area of the spheroid, i.e. with the square of the total radius. This geometric argument leads to the relation:

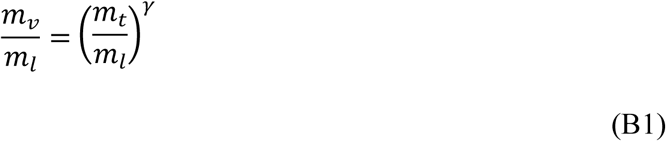

with an asymptotic scaling exponent γ=2/3.

This exponent can also be obtained within the Grimes formalism by applying equation (7) in the limit of large *m_l_*. However, equation (7) further shows that the exact value γ=2/3 is only reached in the theoretical limit of infinitely large tumor spheroids. Experimental observations [4] indicate that the ratio *m_t_/m_l_*, where *m_l_* denotes the spheroid mass at which the central oxygen pressure first vanishes, does not exceed ∼100. Within this biologically relevant range, equation (7) was fitted to the functional form

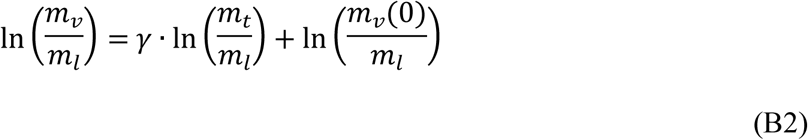

using two separate exponents γ to describe the regimes of small *m_t_/m_l_*(Phase 1) and large *m_t_/m_l_* (Phase 2). The transition point between these regimes was determined via chi-square minimization. The resulting piecewise fit is shown in Figure B1 (dashed line), alongside the viable mass predicted by the Grimes model [5].

For Phase 1 the fit yields γ = 0.835, while in Phase 2 the exponent decreases to γ = 0.713. The maximum deviation between the fitted curve and the Grimes predictions is 0.19% (at *m_t_/m_l_* = 5.5). Notably, the scaling relation (C1) is independent of tumor cell type, as it derives solely from geometric considerations.

---

**Figure B1.**
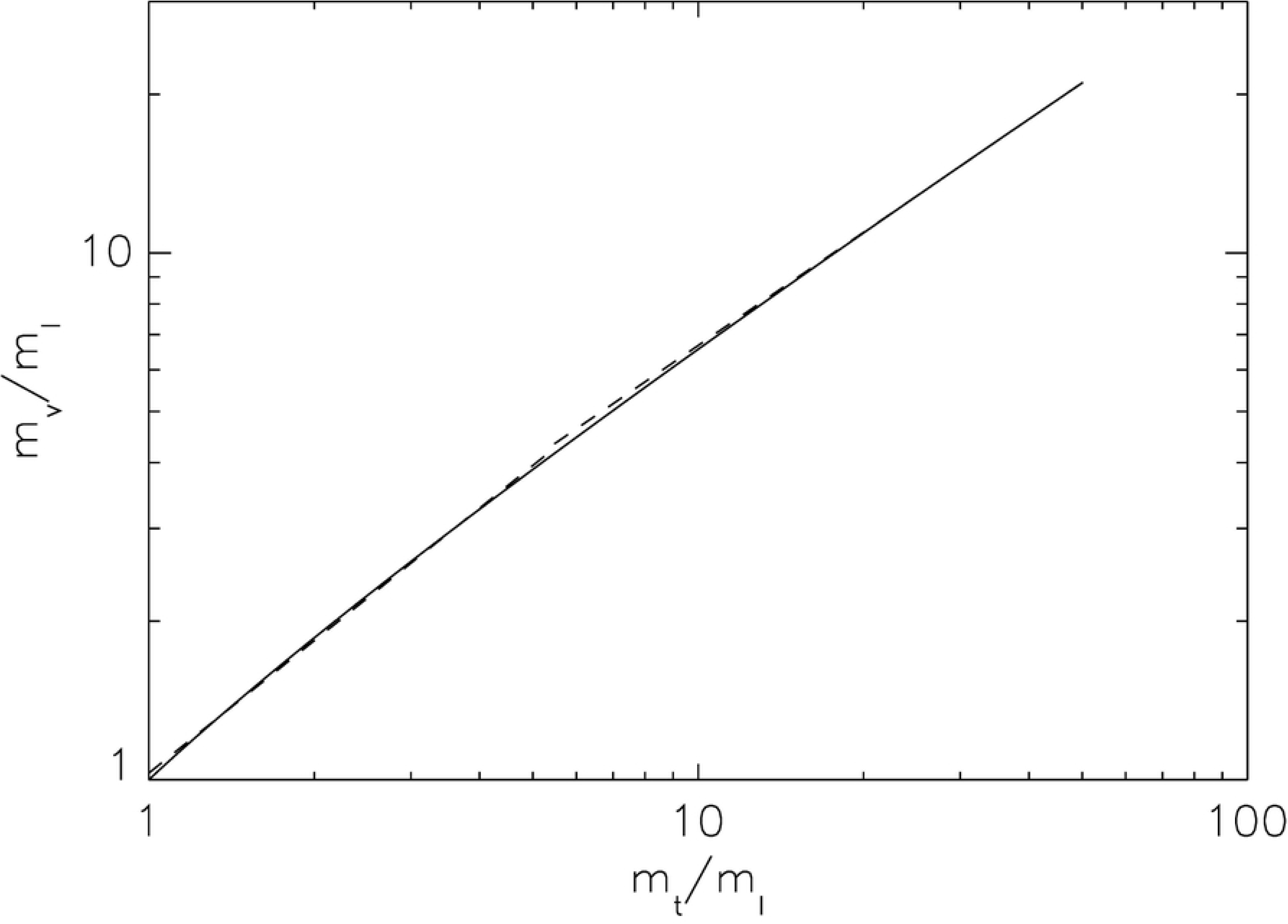
Scaling of viable mass with total spheroid mass. Solid line: Grimes predictions. Dashed line: fitted two-phase scaling relation with exponents *γ* = 0.835 (Phase 1) and *γ* = 0.713 (Phase 2).

## Notes

### Competing Interest Statement

The authors have declared no competing interest.

## References

1. Wallace DI and Guo X (2013) Properties of tumor spheroid growth exhibited by simple mathematical models. Front. Oncol. 3:51. doi: 10.3389/fonc.2013.00051

2. West GB, Brown JH, Enquist BJ (1997) A general model for the origin of allometric scaling laws in biology. Science 276: 122–126.

3. Herman AB, Savage VM, West GB (2011) A Quantitative Theory of Solid Tumor Growth, Metabolic Rate and Vascularization. PLoS ONE 6(9): e22973. doi:10.1371/journal.pone.0022973

4. Freyer, J. P. (1988). Role of necrosis in regulating the growth saturation of multicellular spheroids. Cancer Res. 48, 2432–2439.

5. Grimes DR, Kelly C, Bloch K, Partridge M (2014) A method for estimating the oxygen consumption rate in multicellular tumour spheroids. J. R. Soc. Interface 11: 20131124.

6. Ortega-Arzola E, Higgins PM, Cockell CS (2024) The minimum energy required to build a cell. Scientific Reports 14:5267. 10.1038/s41598-024-54303-6

7. Lynch M, Marinov GK (2015) The bioenergetic costs of a gene. PNAS 112(51) 15690–15695.

8. Marusic M, BajzerZ, Freyer JP, Vuk-Pavlovic S (1994) Analysis of growth of multicellular tumour spheroids by mathematical models. Cell Prolif 27 73–94.

